# Mechanisms for zinc and proton inhibition of the GluN1/GluN2A NMDA receptor

**DOI:** 10.1101/378422

**Authors:** Farzad Jalali-Yazdi, Sandipan Chowdhury, Craig Yoshioka, Eric Gouaux

**Affiliations:** Vollum Institute, Oregon Health and Science University, Portland, Oregon 97239, USA; Department of Biomedical Engineering, Oregon Health and Science University, Portland, Oregon 97239, USA; Howard Hughes Medical Institute, Oregon Health and Science University, Portland, Oregon 97239, USA

## Abstract

*N*-Methyl-D-aspartate receptors (NMDARs) play essential roles in memory formation, neuronal plasticity and brain development with their dysfunction linked to a range of disorders from ischemia to schizophrenia. Zinc and pH are physiological allosteric modulators of NMDARs with GluN2A containing receptors inhibited by nanomolar concentrations of divalent zinc and by excursions to low pH. Despite the widespread importance of zinc and proton modulation of NMDARs, the molecular mechanism by which these ions modulate receptor activity has proven elusive. Here, we use cryo-electron microscopy to elucidate the structure of the GluN1/GluN2A NMDAR in a large ensemble of conformations under a range of physiologically relevant zinc and proton concentrations. We show how zinc binding to the amino terminal domain elicits structural changes that are transduced though the ligand-binding domain and result in constriction of the ion channel gate.

## Introduction

Ionotropic glutamate receptors (iGluRs) are responsible for most of the fast excitatory neurotransmission in the central nervous system and consist of *N*-methyl-D-aspartate (NMDA), kainate, and AMPA receptors (Traynelis et al., 2010). NMDARs are coincidence detectors whose activation requires the binding of glutamate and glycine, and the release of voltage-dependent magnesium block (Nowak et al., 1984). This unique mode of activation coupled with their high calcium permeability underpins the crucial roles of NMDARs in memory formation, the development of long term potentiation, and neuronal plasticity (Bashir et al., 1991; Paoletti et al., 2013). Hyperactivity of NMDARs has been linked to multiple neurological disorders including Alzheimer’s, Huntington’s, and Parkinson’s diseases, whereas their hypoactivity has been linked to cognitive impairment and schizophrenia (Paoletti et al., 2013; Paoletti and Neyton, 2007).

NMDARs are obligate heterotetramers typically consisting of two glycine-binding GluN1 and two glutamate-binding GluN2 subunits (Traynelis et al., 2010). There are eight isoforms of the GluN1 (1a-4b) and four subtypes of GluN2 (A-D). GluN2A and GluN2B subtypes are the most prevalent forms in adult brains, with higher expression levels of GluN2A in the cortex and cerebellum (Akazawa et al., 1994). The first full-length NMDAR (GluN1/GluN2B subtype) crystal structures showed that NMDARs harbor three modular domains: the amino-terminal domain (ATD), the ligand-binding domain (LBD), and the transmembrane domain (TMD) (Karakas and Furukawa, 2014; Lee et al., 2014). The ATDs and the LBDs, comprising the extracellular domain (ECD) of the receptor each adopt clamshell-like structures. The ATD clamshell, in turn, is composed of R1 and R2 lobes, and the LBD clamshell is composed of D1 and D2 lobes. Glycine or glutamate binds at the cleft of the LBD clamshells, resulting in their closure (Furukawa and Gouaux, 2003). A common structural feature of iGluRs is domain swapping, referring to the switching of dimerization partners from the ATD to the LBD (Sobolevsky et al., 2009). In NMDARs extensive interactions between the ATDs and LBDs allows the ATD to allosterically modulate the function of the receptor (Gielen et al., 2009; Paoletti, 2011).

Zinc, protons, polyamines, and ifenprodil are important allosteric modulators of NMDARs. Within a subset of glutamatergic neurons, most notably at the hippocampus, vesicles containing zinc and glutamate are co-released in an activity-dependent manner (Qian and Noebels, 2005, 2006). Zinc-inhibition of NMDARs has a voltage-dependent component, similar to the magnesium block at the receptor pore, and a voltage-independent component (Fayyazuddin et al., 2000). While zinc acts as an allosteric inhibitor of all GluN2 subtypes, GluN2A-containing NMDARs are inhibited by nanomolar concentrations of zinc in a voltage-independent manner, showing a 200-fold greater sensitivity for zinc than GluN2B subunits (Paoletti et al., 1997; Traynelis et al., 1998). Recent studies have discovered evidence of physiological roles for the GluN2A zinc-inhibition. Replacing the wild-type GluN2A subunit with a zinc-insensitive mutant resulted in hypersensitivity to radiant heat or capsaicin and insensitivity to zinc-induced analgesia in mutant mice (Nozaki et al., 2011). Furthermore, two GluN2A mutations identified in patients suffering from childhood-epilepsy and cognitive-deficits have been linked to altered zinc-sensitivity of this subunit (Serraz et al., 2016).

Even though zinc-inhibition is of lower affinity in non-GluN2A subtypes, the extent of inhibition in these subunits is complete, whereas for the GluN2A containing receptors the extent of inhibition is incomplete and dependent on pH (Low et al., 2000). Protons also profoundly modulate NMDAR activity. The open-probability (P_O_) of the receptor is reduced with increasing proton concentration, with a pH IC_50_ value close to the physiological pH (Traynelis and Cull-Candy, 1990). Proton-inhibition is strongly coupled to zinc-inhibition, with studies showing that mutations which increase the sensitivity of the NMDARs to zinc also increase NMDARs’ sensitivity to protons (Fayyazuddin et al., 2000; Gielen et al., 2008). The proton and zinc sensitive sites on NMDARs were mapped to the ATD by a series of electrophysiological studies (Fayyazuddin et al., 2000; Gielen et al., 2009; Low et al., 2000; Paoletti et al., 2000). Subsequent structures for GluN2A and GluN2B isolated-ATDs unequivocally demonstrated the high-affinity zinc-binding site and the coordination of zinc by four amino acid sidechains (H44 and H128 in the R1 lobe, E266 and D282 in R2 lobe) (Karakas et al., 2009; Romero-Hernandez et al., 2016).

While functional and electrophysiological studies have pinpointed domains involved in zinc and proton modulation of NMDA receptor activity, the molecular mechanism for this inhibition is unresolved. On the one hand, Gielen et al. showed that enhancing the LBD heterodimer stability reduced the extent of zinc-inhibition, whereas disrupting this interface led to more potent inhibition, establishing an important role for the D1-D1 interface in zinc-inhibition (Gielen et al., 2008). On the other hand, luminescence resonance energy transfer (LRET) studies point toward a conserved structural mechanism for zinc and ifenprodil inhibition (Sirrieh et al., 2013, 2015), and the structures of the NMDA receptor bound to ifenprodil or other phenylethanolamines revealed an intact D1-D1 LBD interface (Tajima et al., 2016; Zhu et al., 2016). Consistent with the view of a conserved structural mechanism of zinc and ifenprodil inhibition, ifenprodil enhances zinc and proton sensitivity (Amico-Ruvio et al., 2012), and in the case of GluN1/GluN2A/GluN2B triheteromeric receptors, zinc enhances ifenprodil sensitivity and the extent of inhibition (Hansen et al., 2014). Finally, single-channel recordings and single-molecule FRET (smFRET) studies point towards an ensemble of electrophysiologically silent states, where the NMDA receptor primarily populates the pre-open or low P_O_ states in the presence of zinc (Amico-Ruvio et al., 2011; Dolino et al., 2017).

In this work we solve the first full-length structure of the GluN1/GluN2A diheteromeric receptor and elucidate the structural underpinnings of proton-dependent zinc-inhibition for the GluN2A-containing NMDARs via single-particle cryoEM. We demonstrate closure of the ATD clamshells due to zinc-binding and the transduction of this conformational change through the LBD and finally to the pore. The seven distinct structural conformations observed during the course of this study, some with an intact D1-D1 interface and others with a disrupted D1-D1 interface, illustrate how zinc and proton inhibition leads to a lessening of the tension in the linkers connecting the LBDs to the channel pore, leading to constriction of the ion channel gate.

## Results and Discussion

### Function of the GluN1/GluN2A receptor in membranes and micelles

The GluN1/GluN2A diheteromeric receptor (diNMDAR) construct is truncated to exclude the largely unstructured C-terminal domains (CTD) of GluN1 and GluN2A after residues 847 and 866, respectively. To investigate the zinc and proton modulation of the diNMDAR construct, we performed two electrode voltage clamp (TEVC) experiments, using buffered zinc solutions (Paoletti et al., 1997). Divalent zinc inhibits diNMDAR currents with an IC_50_ of 12 nM at pH 6.8, and 36 nM at pH 7.4 (Figure 1A and S7A-B). The magnitude of voltage-independent maximal zinc-inhibition was ~50% at pH 7.4 and ~80% at pH 6.8, in agreement with previously measured values (Low et al., 2000).

**Figure 1.**
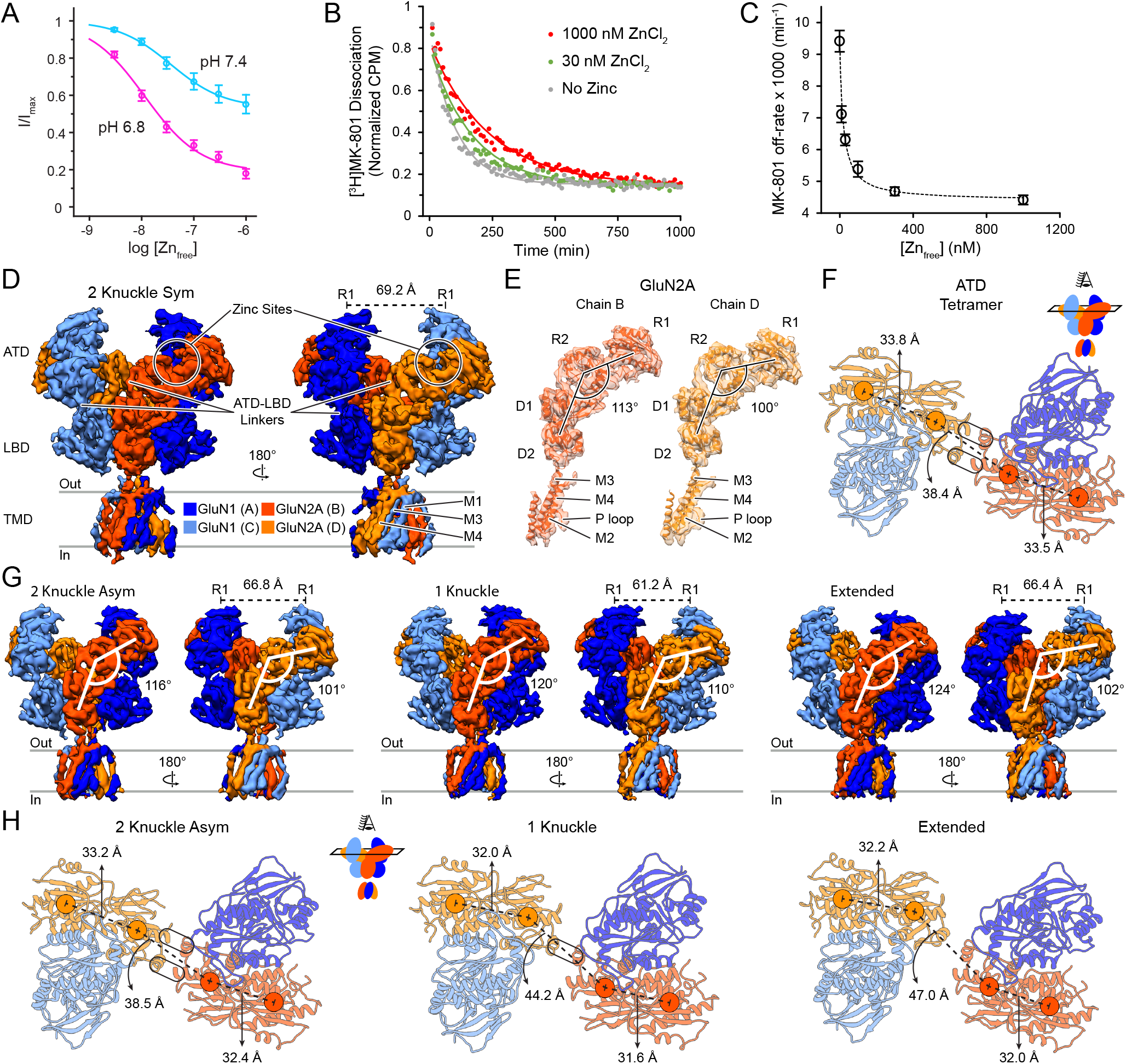
GluN1/GluN2A Receptor Activity Modulation by Proton and Zinc, and CryoEM Structures at pH 7.4 in the Absence or Presence of 1 μM Zinc. (A) Zinc-inhibition of the diNMDAR construct at pH 7.4 (cyan, IC_50_ = 36 ± 1 nM) and pH 6.8 (magenta, IC_50_ = 12 ± 1 nM) in the presence of 100 μM glutamate and glycine. Imax is obtained by measuring currents in the absence of zinc. Data are mean ± SEM of 3-4 oocytes. (B) Dissociation of [^3^H]MK-801 from agonist-bound affinity-purified diNMDAR (100 μM glutamate and glycine) in the absence of zinc (grey), in the presence of 30 nM zinc (green), or 1 μM zinc (red). A single-exponential curve (solid lines) is used to obtain the off-rate of MK-801. (C) Off-rate of [^3^H]MK-801 as a function of zinc at pH 7.4 in the presence of agonists (100 μM glutamate and glycine). Zinc-inhibition IC_50_ = 17.3 ± 1.4 nM based on a one-site binding model. Data are mean ± SEM of three independent replicates. (D and G) Side views of the diNMDAR cryoEM map in the presence of agonists (1 mM glutamate and glycine) at pH 7.4 with 1 mM EDTA (D) or in the presence of 1 μM zinc (G). The distance between the center of mass (COM) of the upper lobes (R1) of GluN1 ATD are shown. (E) Atomic model of each GluN2A subunit fitted into the density map. The ATD-LBD angle, angle between a vector passing through the COM of R1 and R2 lobes of ATD and a vector passing through the COM of D1 and D2 lobes of LBD, is indicated for the two subunits. (F and H) Top view of the atomic model for the ATD of the diNMDAR in the absence of zinc (F) or in the presence of 1 μM zinc (H). COMs of R1 and R2 lobes for the two GluN2A subunits are shown (circles), with their distance indicated. The rounded rectangles represent the ‘knuckles’ and show the interaction between the α5 and α6 helices of opposing GluN2A subunits for the 2-knuckle conformations, and between α6 and α6 helices for the 1-knuckle conformation.

To interrogate zinc modulation of receptor ion channel gating in detergent micelles, the conditions under which the cryoEM studies are performed, we designed a radiolabeled binding assay using the channel blocker MK-801. MK-801 is an NMDA receptor ion channel blocker whose binding and un-binding rates increase when the receptor is in the open conformation (Blanke and VanDongen, 2008; Kornhuber et al., 1989; Song et al., 2018). Using a scintillation proximity assay we measured the off-rate of [^3^H]MK-801 from receptors in detergent micelles in the presence of agonists and at various concentrations of zinc (Figure 1B). In the presence of saturating zinc, the off-rate of MK-801 is slowed ~2-fold at pH 7.4. Using a single-site binding model, the off-rate of MK-801 as a function of free-zinc concentration has an estimated IC_50_ of ~20 nM (Figure 1C). The similarity between zinc-inhibition IC_50_ and the extent of inhibition obtained via electrophysiology and radiolabeled binding assays indicate that zinc-modulation of the receptor is retained for the diNMDAR construct used in these studies, in detergent micelles.

### Architecture of the GluN1/GluN2A receptor in the absence of zinc

We solved the structure of the GluN1/GluN2A receptor in the presence of agonists and 1 mM EDTA using single-particle cryoEM at 6.2 Å resolution (Figure 1D and S1D-F) and modeled each subunit’s ATD and LBD lobes, as well as the TMD, into the density by rigid-body fitting (Figure 1E). The homology model for the GluN1 subunit was based on the GluN1/GluN2B crystal structure (PDB code: 4PE5) (Karakas and Furukawa, 2014) and the model for the GluN2A subunit was based on the crystal structure of the isolated GluN1/GluN2A ATD (Romero-Hernandez et al., 2016) (PDB code: 5TQ0) and the isolated LBD of GluN1/GluN2A (PDB code: 5I57) (Yi et al., 2016), with the remainder of the homology model being based on the cryoEM structure of the GluN1/GluN2A/GluN2B triheteromeric NMDAR (PDB code: 5UOW) (Lu et al., 2017).

The overall architecture of the receptor resembles a bouquet with layered ATD, LBD, and TMD segments, as seen in other subtypes of the NMDA receptor (Karakas and Furukawa, 2014; Lee et al., 2014; Lu et al., 2017). Similar to the GluN1/GluN2B receptor (Zhu et al., 2016), the ion channel gate in the presence of agonists and in the absence of zinc is closed, thus we speculate that the receptor is in the desensitized or pre-open conformation. The receptor exhibits pseudo C2-symmetry along the central axis of the channel. At the ATD, the R2 lobes of opposing GluN2A subunits meet at the pseudo-symmetry axis, with the N-terminus of the α5 helix interacting with the α6 helix of the opposing subunit (Figure 1F). Viewed from ‘above’, the R2 lobes of GluN2A subunits resemble two hands performing a ‘fist bump’ with two of the knuckles (α5 and α6) in each fist meeting two knuckles from the opposing fist. Due to the overall pseudosymmetry and fist-bump resemblance with 2 contacting knuckles, we named this conformation 2-knuckle-sym. In this conformation, the GluN2A ATD clamshells are in the ‘open’ configuration, with R1-R2 center of mass (COM) distance between 33.5-33.8 Å (Figure 1F).

A striking deviation from the pseudo-C2-symmetry in the 2-knuckle-sym conformation is the angle between the ATD and LBD of the GluN2A subunits. When we measured the angle between two vectors, the first of which passes through the COMs of the LBD D1 and D2 lobes, and the second of which passes through COMs of the ATD R1 and R2 lobes, defined as the ATD-LBD angle, we noticed a 13° difference with respect to the two GluN2A subunits (Figure 1E). This deviation from symmetry can also be measured in the lengths of GluN2A LBD-TMD linkers and the dimer-dimer distances at the LBD. We used the ATD-LBD angle to assign chain B (the larger angle) and chain D (the smaller angle) for all of the structures in this study.

### An ensemble of conformations at nanomolar zinc concentration

To elucidate the structural basis for nanomolar inhibition of ion channel activity by zinc, we dialyzed the receptor against a buffer containing glutamate, glycine and 1 μM ZnCl_2_ prior to EM grid preparation. We collected ~1,600 micrographs, resulting in ~270,000 particles after ‘clean up’ by 2D-classification which were then used for 3D-classification. The resolution of the observed three classes is between 6.1 – 7.1 Å, allowing us to independently rigid-body fit structures of each subunit’s ATD and LBD lobes, as well as the TMD into the density (Figure S2).

The class that most closely resembles the ‘EDTA’ conformation (Figure 1G-H, left) adopts a ‘2-knuckle’ structure and accounts for 19% of the particles. The most noticeable difference between this class and the ‘EDTA’ conformation is the state of the ATD clamshells. Here, the R1 lobe of GluN2A chain B moves 4.6 Å with respect to the 2-knuckle-sym structure, closing the ATD clamshell by 10.3°, and reducing the R1-R2 COM distances by 1.1 Å. The closure of the ATD clamshell in chain B is accompanied by a 3° increase in the ATD-LBD angle for this chain with respect to the 2-knuckle-sym conformation. Changes between the ‘EDTA’ conformation and the current class are much smaller for chain D, where the change in ATD-LBD angle remains indistinguishable at the limit of current resolution, accompanied by the R1-lobe movement of 1.1 Å, closure of the ATD clamshell of 2.8°, and decrease in the R1-R2 COM distance of 0.6 Å. The position of the opposing GluN2A R2 lobes at the pseudosymmetry axis is unaltered in this class, and indeed the COM distance between the opposing R2-R2 lobes remains similar to the 2-knuckle-sym conformation. Due to the asymmetry that is now present with regard to the ATD clamshell closures, we named this structural class the 2-knuckle-asym conformation.

In the second class, accounting for 24% of the particles (Figure 1G-H, middle), R2 lobes at the pseudo-symmetry axis have moved so only the opposing α6 helices are poised for subunit-subunit interaction. This is the ‘1-knuckle’ class, as it has only one point of contact between the two ATD ‘fists’. The movement of the R2 lobes increases their COM distance by 5.8 Å and is accompanied by the closure of both ATD clamshells. For chain B, the subunit with the larger ATD-LBD angle, the R1 lobe moves by 6.1 Å leading to an ATD clamshell closure of 11.9°, and bringing the R1 and R2 lobes closer together by 1.9 Å as compared to the 2-knuckle-sym conformation. Similarly, for chain D, the R1 lobe moves by 5.8 Å to close the ATD clamshell by 15.4°, bringing the R1 and R2 COMs closer by 1.8 Å. Along with the ATD closure, there is an increase in the ATD-LBD angle for chains B and D by +7° and +10°, respectively.

The third class accounts for 17% of the particles. In this class, one of the ATD dimers (chains C and D) moves away from the pseudo symmetry-axis while the other ATD dimer remains in place. The movement of one ATD dimer away from the rest of the receptor results in no interaction between the R2-lobes of GluN2A subunits and an increase in the distance between the COM of Chain D’s R2 to D1 lobes of 8.4 Å (Figure 1G-H, right). We named this class the ‘extended’ conformation due to the extension of one ATD dimer from the rest of the receptor. Both GluN2A ATD clamshells are closed with the R1-R2 COM distance being reduced by ~1.6 Å for each chain with respect to the 2-knuckle-sym conformation. We also noticed that the extended ATD dimer shows more diffuse density as compared to the rest of the ECD, suggestive of conformational heterogeneity. The ATD-LBD angle in chain B is 4° greater in the extended conformation compared to the 1-knuckle conformation, however, the angle chain D is 102°, a value similar to the 2-knuckle conformations.

### Conformations of the GluN1/GluN2A/GluN2A* triheteromeric receptor

The 2-knuckle-asym conformation is consistent with one ATD dimer in a zinc-bound, closed clamshell conformation and the other indistinguishable from the open clamshell conformation. To investigate whether the closed and open conformations are the consequence of zinc-bound states, and to probe the extent to which conformational changes are propagated from one ATD heterodimer throughout the receptor, we carried out cryoEM studies on a ‘triheteromeric’ GluN1/GluN2A/GluN2A* receptor where the GluN2A* subunit harbors a His 128 to Ser mutant. This substitution, at the high affinity GluN2A ATD zinc site, decreases zinc potency by ~500-fold and reduces the extent of zinc inhibition (Fayyazuddin et al., 2000; Hansen et al., 2014; Hatton and Paoletti, 2005; Paoletti et al., 2000). To distinguish the GluN2A and GluN2A* subunits in cryoEM density maps, we knocked out a prominent glycosylation site (N687) in the LBD layer of the His 128 to Ser GluN2A* subunit.

We first solved the structure of the GluN1/GluN2A/GluN2A* triNMDAR in the presence of agonists and 1 mM EDTA at 5.4 Å resolution. This structure (Figure 2A and Figure 1S) is similar to the 2-knuckle-sym conformation of the GluN1/GluN2A diNMDAR, showing the same asymmetry with respect to the ATD-LBD angle for the two GluN2A subunits. For the triNMDAR in the absence of zinc, we do not observe a signal consistent with glycosylation at either subunit, whereas for the diNMDAR the glycosylation density is substantial for both subunits (Figure 2B-C). The lack of glycosylation density is likely due to the averaging of the density for the GluN2A and GluN2A* at each position for the triNMDAR, as both subunits are expected to behave identically in the absence of zinc.

**Figure 2.**
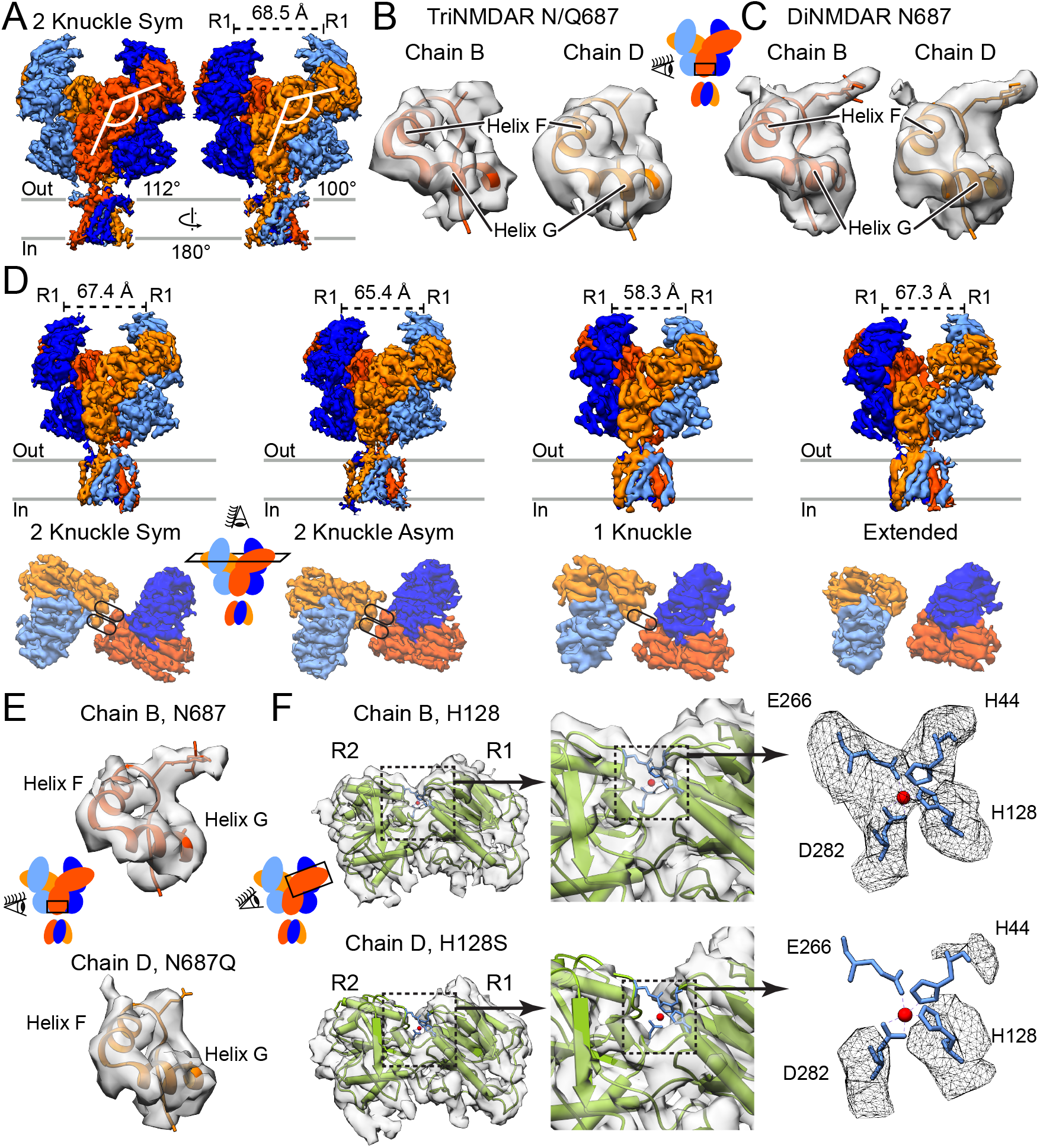
CryoEM Structures of the GluN1/GluN2A/GluN2A* triNMDAR. (A) Side views of the GluN1/GluN2A/GluN2A* triNMDAR cryoEM map in the presence of agonists (1 mM glutamate and glycine) at pH 7.4 with 1 mM EDTA. The distance between COM of the upper lobes (R1) of GluN1 ATD, as well as the ATD-LBD angle for the two GluN2A subunits, are shown. (B and C) Atomic model and the associated map for the GluN2 subunit of the triNMDAR (B) and the diNMDAR (C) in the vicinity of residue 687. There is density consistent with glycosylation of N687 in the diNMDAR, however this density is not present in either subunit for the triNMDAR. Since distinguishing between GluN2A and GluN2A* subunits was not possible based on glycosylation at residue 687, residue 687 in the map was represented as an alanine. (D) Side view of the full receptor (top) and top view of the ATD (bottom) of the GluN1/GluN2A/GluN2A* triNMDAR cryoEM map in the presence of agonists (1 mM glutamate and glycine) and 1 μM zinc at pH 7.4. (E) Atomic model and the associated map for the GluN2A subunit (Chain B, top) and the GluN2A* subunit (Chain D, bottom) in the vicinity of residue 687 for the 2-knuckle-asym conformation of the triNMDAR in the presence of 1 μM zinc. At the same contour level, there is density consistent with glycosylation at one GluN2 subunit and not the other, allowing for the identification of the wt GluN2A and the H128S & N687Q mutant GluN2A subunit. (F) The crystal structure of the GluN2A ATD bound by zinc (PDB: 5TPW) fitted into the cryoEM map for the R1 lobe of the two GluN2 subunits of the triNMDAR 2-knuckle-asym conformation. There is density consistent with four amino acid sidechains (H44, H128, E266, D282) coordinating zinc at the zinc-binding site for the WT-subunit. At the same contour level, there is no density in the vicinity of the zinc-binding pocket for the GluN2A* subunit with the H128S mutation.

In the presence of 1 μM zinc, the triNMDAR occupies the four conformations that we had observed previously, with resolutions ranging from 5.3 Å for the 2-knuckle-asym to 7.5 Å for the extended conformation (Figure 2D-E and S3). Compared to the diNMDAR, the triNMDAR particles occupied a larger fraction in the 2-knuckle classes (52% in triNMDARs, 19% in diNMDARs) than the 1-knuckle conformation (13% in triNMDARs, 24% in diNMDARs). The extended conformation accounts for ~17% of the particles for both constructs. To further increase the resolution of the 2-knuckle-asym class we performed a focused-refinement on the ECD portion of the channel, which allowed us to improve the resolution to ~4.7 Å (Figure S7C-L). The structures for ECD lobes and TMD segment of each subunit was rigid-body fitted into the maps as described above.

The 2-knuckle-asym class comprised the largest fraction of the particles among observed conformations (28%) and presented the most clear distinction between the GluN2A (chain B) and GluN2A* (chain D) subunits based on the glycosylation density at N687 (Figure 2F). Here, chain B adopts a closed ATD clamshell conformation with a larger ATD-LBD angle, and chain D adopts an open clamshell conformation. We next rigid-body fitted the x-ray structure of the isolated GluN2A ATD bound by zinc (Romero-Hernandez et al., 2016) into both GluN2A subunit’s R1 lobe density. There is density associated with the zinc-binding pocket in chain B (Figure 2G). This density has a tetrahedral shape and is consistent with zinc being coordinated by sidechains of H44, H128, E266, and D282 residues. By contrast, there is no density around the zinc ion when the model is fitted into the density for chain D, consistent with GluN2A*’s lack of zinc-sensitivity. With the triNMDAR 2-knuckle-asym cryoEM map, representing the highest resolution map obtained in this study, we were able to confirm that the 2-knuckle-asym class represents dimorphic GluN2A ATDs, with one in the open, apo zinc state and the other in the closed, zinc bound conformation.

The triNMDAR 1-knuckle and extended conformations exhibited both ATD clamshells in the closed conformations, even though one subunit lacks the high-affinity zinc-binding site. It is highly unlikely that closed ATD clamshells in the 1-knuckle and extended conformations of triNMDAR represent the GluN2A diheteromeric contaminant in the triNMDAR sample, as the density at the N687 residue consistent with glycosylation which was prominent in the GluN2A diheteromeric receptors is absent in these structures. The closed ATD clamshells of both subunits in these conformations is consistent with our findings that the ATD conformation does not represent an isolated structural motif, and that conformational changes in other domains can induce alternative ATD clamshell conformations.

### Conformational changes due to proton and millimolar zinc concentration

Electrophysiology experiments show that the extent of zinc inhibition is ~98% at pH 6.2 (Figure S7B). To understand how protons enhance the extent of zinc inhibition, we prepared the GluN1/GluN2A receptor in a MES buffer at pH 6.1 in the presence of agonists and 1 μM zinc. Analysis of 2D class averages shows a number of 2D classes with the ATD dimers farther apart than observed previously (Figure 3A). After 3D-classification, ~9% of the particles belong in a structural class where the two dimers have lost the ATD to the LBD domain swapping and show a ‘super splayed-open’ conformation (Figure 3B, right). The splayed conformation also shows a disruption of the D1-D1 interface within the LBD dimers. Other than the super-splayed class (Figure S4I-J), ~30% of the particles are classified into the 1-knuckle conformation (Figure S4C-E), and ~25% of the particles are classified into the extended conformation (Figure S4F-H). We did not observe any classes which show a 2-knuckle conformation. For the 1-knuckle and the extended conformations, the resolution of the maps was sufficiently high to allow for rigid-body fitting of each subunit’s individual ECD lobes and the TMD into the density. For the super-splayed conformation, each ATD heterodimer, as well as the TMD tetramer, had to be rigid-body fit as a single unit. The LBD of the receptor was then rigid-body fit as a single domain per subunit.

**Figure 3.**
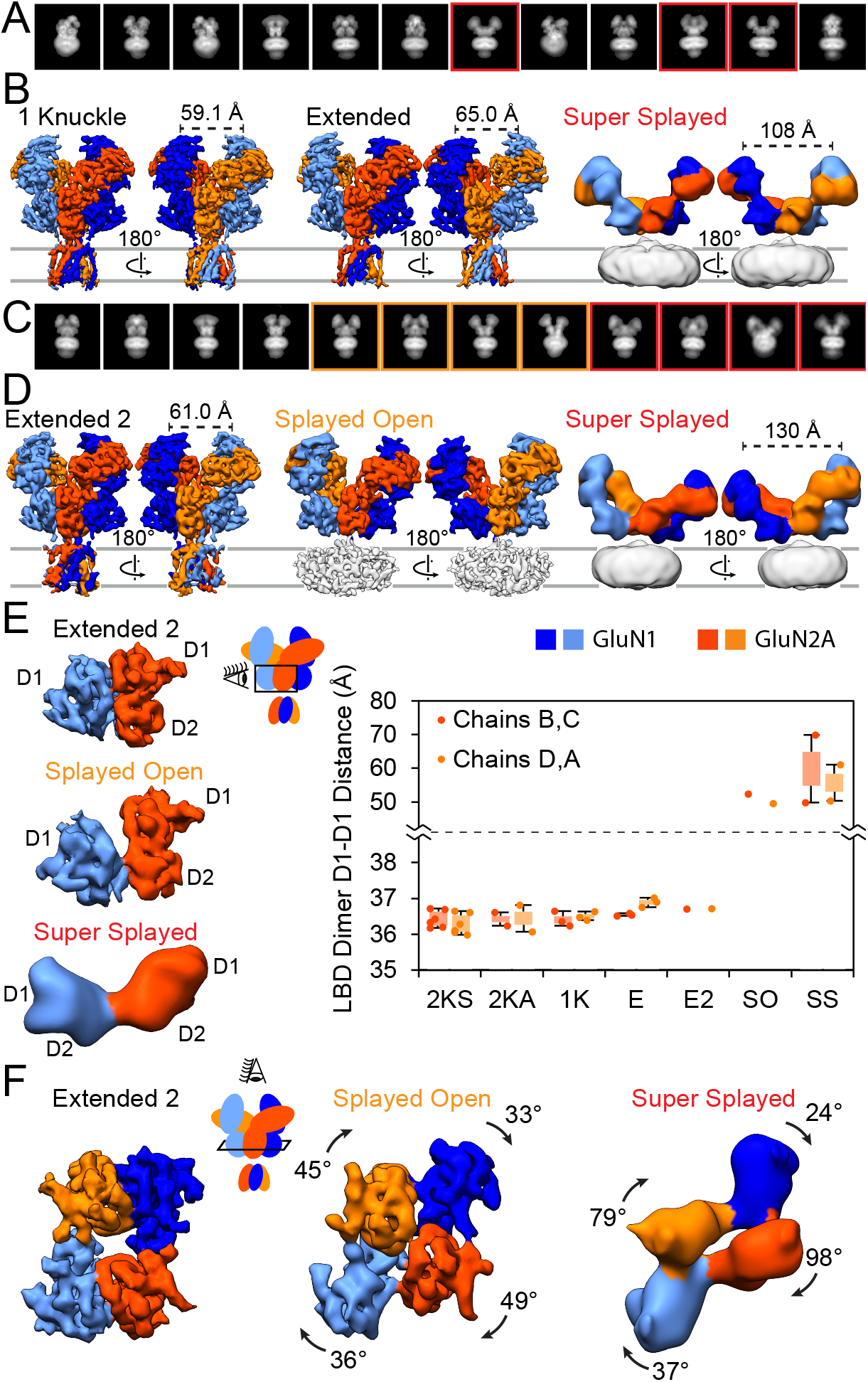
CryoEM Structures of the GluN1/GluN2A diNMDAR in High Proton and Zinc Concentrations. (A) Representative 2D class averages of the diNMDAR in the presence of agonists (1 mM glutamate and glycine) and 1 μM zinc at pH 6.1, with red boxes outlining the super-splayed conformation. (B) Side views of the diNMDAR cryoEM map in the presence of agonists (1 mM glutamate and glycine) and 1 μM zinc at pH 6.1. (C) Representative 2D class averages of the diNMDAR in the presence of agonists (1 mM glutamate and glycine) and 1 mM zinc at pH 7.4, with orange and red boxes outlining the splayed and the super-splayed conformations, respectively. (D) Side views of the diNMDAR cryoEM map in the presence of agonists (1 mM glutamate and glycine) and 1 mM zinc at pH 7.4. (E) The intact LBD dimer of the extended-2 conformation (top) breaking apart in the splayed (middle) and the super-splayed (bottom) conformations, with the GluN1 and GluN2A D1 lobes losing their interactions. The graph on the right shows the distance between the COM of the D1 lobe of GluN1 to the D1 lobe of GluN2A at the major interface for the seven structural classes (2KS = 2-knuckle-sym, 2KA = 2-knuckle-asym, 1K = 1-knuckle, E = extended, E2 = extended-2, SO = splayed-open, SS = super-splayed). Bars represent the first and third quartile of each population, and the error bars represent the maximum and minimum values. (F) Top view of the LBD for the structures in (D). There is a clockwise rotation for each subunit’s LBD when the D1-D1 interface is broken in the splayed-open or super-splayed conformations as compared to the extended-2 conformation.

As a counterpoint to decreasing the pH to obtain maximal zinc-inhibition, we examined the structure of the receptor at high pH in the presence or absence of zinc to assess conformational changes leading to minimal zinc-inhibition. We acquired data sets of the GluN1/GluN2A receptor at pH 8.0 in 0.1 mM EDTA, or at pH 8.0 and 1 μM zinc (Figure S6). Both datasets only yielded the 2-knuckle-sym conformation after 3D-classification.

We next examined the conformations of the receptor in the presence of 1 mM zinc at pH 7.4. The voltage-independent component of zinc-inhibition plateaus at ~1 μM zinc, and further increasing the zinc concentration does not lead to more voltage-independent inhibition (Paoletti et al., 1997). There exist, however, additional zinc-binding sites on GluN1 and GluN2B ATDs (Karakas et al., 2009; Romero-Hernandez et al., 2016), and we were interested to learn if there are distinct structural classes associated with high zinc concentrations. Indeed, there are glutamatergic neurons with millimolar zinc concentrations inside presynaptic vesicles (Sensi et al., 2009) and upon release synaptic zinc concentrations can be greater than 100 μM (Vergnano et al., 2014).

We acquired a dataset with the GluN1/GluN2A receptor at pH 7.4 in the presence of agonists and 1 mM zinc. It was clear from the micrographs and 2D-class averages that many of the receptor particles have adopted splayed-open conformations, with their ATD dimers far away from each other (Figure 3C). After 3D-classification, three distinct classes emerged (Figure 3D and S5). The first class (46% of the 2D-cleaned up particles) shows similar structural elements to the extended classes we had observed previously, and was named the extended-2 class. However, the degree of separation between the COMs of R2 lobes of GluN2A subunits in the extended-2 class is 5.7 Å larger than the extended class. Another difference between the extended-2 and the extended conformations is the distance between the COMs of GluN2A R2 and D1 lobes for the extended ATD dimer, which is ~5.3 Å lower in the extended-2 conformation.

The second class we encountered at 1 mM zinc concentration shows a disrupted D1-D1 interface at the LBD dimers with moderately separated ATD dimers, and was thus labeled the splayed-open class. While this class originally accounted for ~16% of the 2D-cleaned up particles, the resolution of the class was improved by subsequent rounds of 3D-classification and refinement. For the splayed-open class, we were able to rigid-body fit each lobe of the ECD into the density independently. However, due to the poor resolution at the TMD, the TMD tetramer had to be fitted as a single body (as described in methods). This class represents the highest resolution structure of the NMDA receptor in a conformation with a disrupted D1-D1 interface. The final class that we observed shows a similar conformation to the super-splayed class seen in the low-pH dataset, and accounted for ~15% of the 2D-cleaned up particles before additional rounds of 3D-classification and refinement to improve the resolution.

To confirm that the conformations in the presence of 1 mM zinc did not represent unfolded receptor due to the millimolar zinc concentrations, after the initial grids were frozen with 1 mM zinc we added 3 mM EDTA to the remainder of the sample to chelate zinc and prepared cryoEM samples. After 2D-cleanup of the particles, 3D-classification and refinement led only to the 2-knuckle-sym conformation (Figure S1J-M). Neither during the 2D-nor 3D-classification did we observe any classes which represented the splayed open conformations. These results are consistent with the splayed-open or the super-splayed conformations being reversible structures which can transition to other conformations when zinc is removed from the solution.

We analyzed the LBD dimer D1-D1 separation (COM distance) for all of the structures that we had solved (Figure 3E). We did not observe any classes other than the splayed-open and super-splayed where the D1-D1 interface was altered substantially. The super-splayed conformation has been observed before in the context of the NMDARs. Zhu et al., demonstrated that the GluN1/GluN2B NMDAR adopts a spectrum of different conformations when bound by competitive antagonists, five of which showed splayed ATD dimers and disrupted D1-D1 interfaces (Zhu et al., 2016). One of the characteristics of GluN1/GluN2B splayed classes was the clockwise rotation of the GluN2B LBD clamshells by 92-118°. For the GluN2A LBDs, we observed a 45-49° clockwise rotation in our splayed-open class, and a 79-98° rotation in the super-splayed class as compared to the extended-2 conformation (Figure 3F).

### Zinc-induced conformational rearrangements at the ATD

The GluN2A clamshell undergoes closure upon zinc-binding. For the 2-knuckle-sym, 2-knuckle-asym, 1-knucle, and extended conformations, the resolution of the individual structures was sufficient to allow rigid-body fitting the structure of individual lobes into the density. For these conformations, we measured the R1-R2 COM distance for the two different GluN2A subunits. The open clamshell is centered at ~33.6 Å of R1-R2 separation, whereas the closed clamshell is centered at ~32.0 Å of separation (Figure 4A). While chains B and D in the 2-knuckle-sym class are in the open ATD clamshell conformation, only chain B adopts the closed clamshell conformation as the receptor transitions to the 2-knuckle-asym conformation.

**Figure 4.**
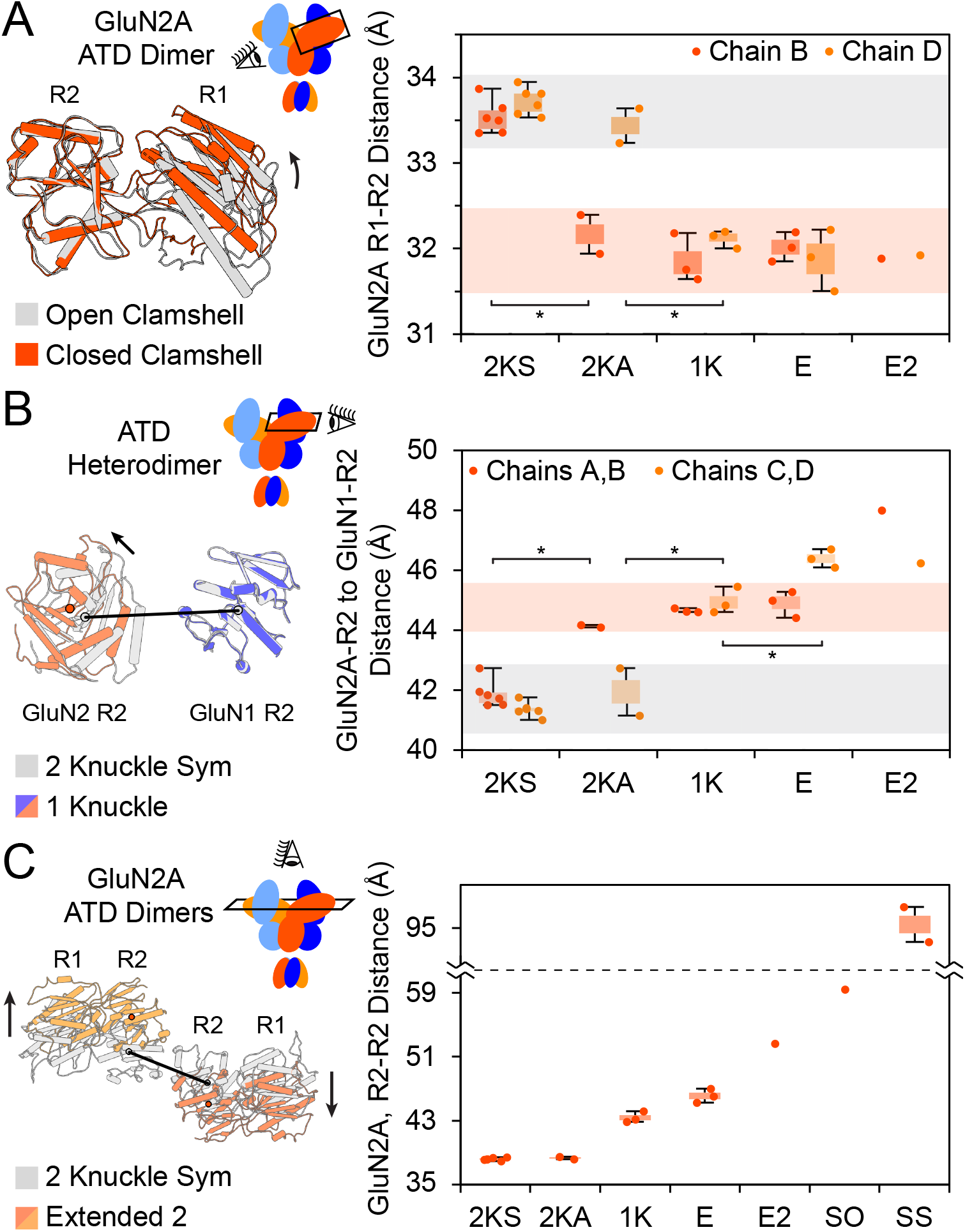
Zinc-Induced Conformational Rearrangements at the ATD. (A) Distance between COMs of the R1 and R2 lobes of GluN2A subunits for the various structural classes. The area shaded in grey represents the open ATD clamshell, and the area shaded in red represents the closed ATD clamshell. The extended-2, splayed-open and super-splayed conformations were not included in this analysis due to poor resolution. Side view of the GluN2A ATD in the closed and open clamshell conformations is shown to the right, with the R2 lobes aligned. The black arrow indicates the direction of motion. * denotes significance (p < 0.05) using a two-tailed homoscedastic student’s t-tests. (B) Closure of the ATD clamshell increases the distance between the GluN1 and GluN2A R2 lobes. The grey shaded area represents the open GluN2A ATD clamshell, and the red shaded area represents the closed GluN2A ATD clamshells with close ATD-LBD association. Side view of the GluN1 and GluN2A R2 lobes in the closed and open GluN2A clamshell conformations is shown to the right, with the R2 lobes of GluN1 aligned. (C) When both ATD clamshells are closed (1K-SS), the GluN2A R2 lobes move away from each other perpendicular to the R1-R2 axis. Top view of the GluN2A ATDs in the 2-knuckle and extended-2 conformations is shown to the right, aligned by their pseudosymmetry axes. The black arrows indicates the direction of motion.

The R2 lobes of GluN1 and GluN2A ATD play a critical role in transduction of ATD conformational changes to the LBD, as they are directly connected to the LBD D1 lobes. Upon zinc-binding and the closure of the GluN2A ATD clamshell, the R2 lobes of GluN1 and GluN2A within each ATD heterodimer move apart by ~3 Å (Figure 4B). Chain D of the extended conformation experiences an additional 1.4 Å expansion (p = 0.01) when the ATD heterodimer extends from the receptor and loses interaction with the LBD layer.

Movement of the R2 lobes of GluN2A subunits at the pseudo-symmetry axis is also associated with zinc and proton inhibition of the receptor. In all classes with an intact D1-D1 interface at the LBD dimer, the R2 lobes of GluN2A move perpendicular to the R1-R2 axis, resulting in lateral movement of the lobes away from each other and the formation of the 1-knuckle and extended conformations (Figure 4C). This lateral R2-R2 movement is only observed in conformations with both ATD clamshells closed.

### Transduction of conformation changes from the ATD to the LBD

To understand how conformational changes within the ATD layer are transduced to the LBD layer, we first examined the ATD-LBD angle. When the ATD clamshell of chain B closes, as the receptor transitions from the 2-knuckle-sym to the 2-knuckle-asym conformation, the ATD-LBD angle for this chain increases by ~6° (Figure 5A). In contrast, the ATD-LBD angle remains unchanged for chain D, which does not undergo ATD clamshell closure. In the 1-knuckle conformation, where both ATD clamshells are closed, there is a 12° increase in the chain D ATD-LBD angle and a 6° increase for chain B as compared to the 2-knuckle-sym state. In the case of chain D of the extended conformation, there is an uncoupling of the ATD clamshell state from the rest of the receptor, where the ATD clamshell is closed even though the angle for chain D is ~101°, the ATD-LBD angle of the open clamshell state. The increase of the ATD-LBD angle upon closure of the ATD clamshell is a direct result of the R1-lobe of the GluN2A ATD moving to close the clamshell, with the R2 lobes remaining at the central axis of symmetry.

**Figure 5.**
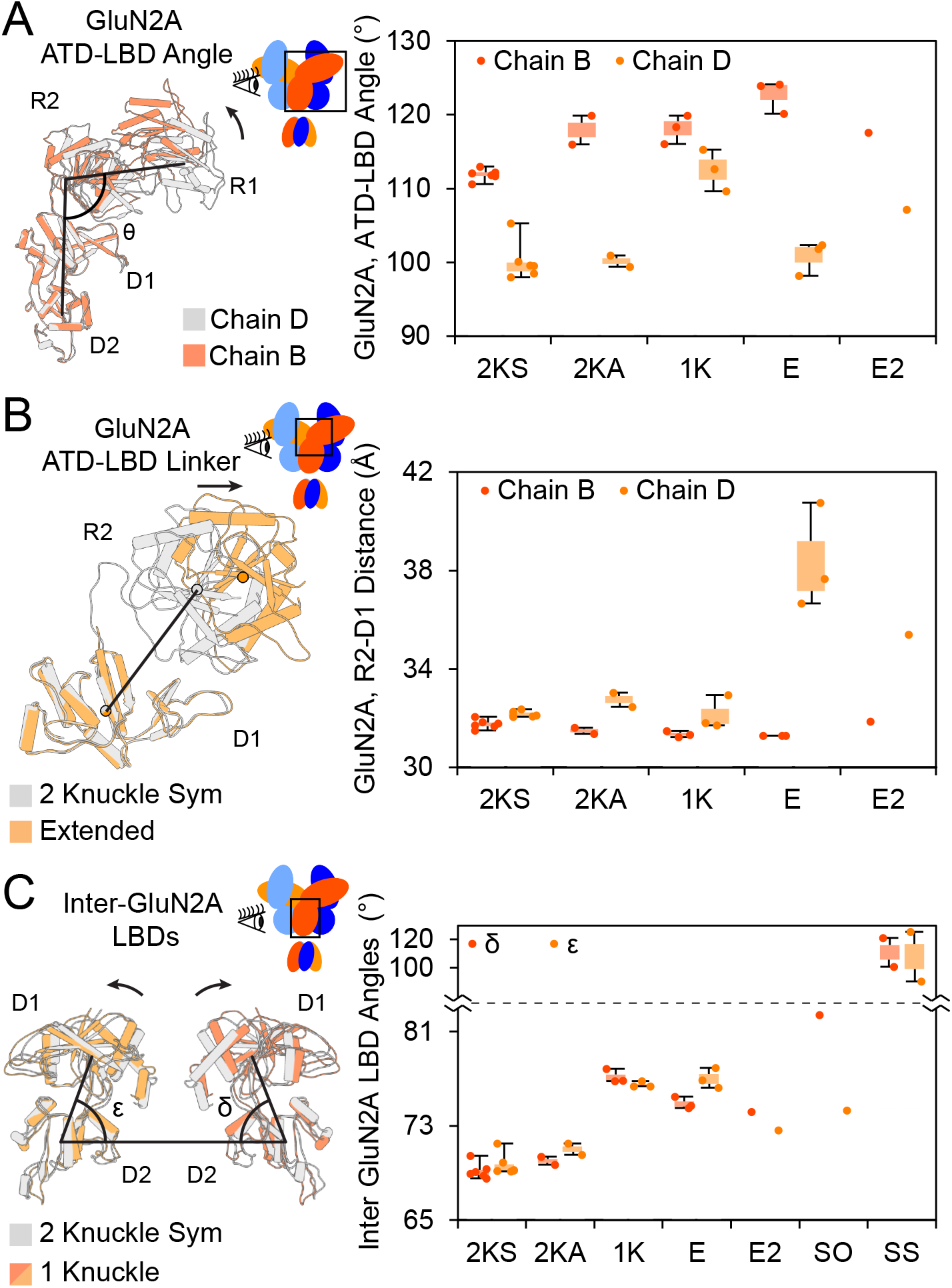
Translation of Conformation Changes from ATD to the LBD. (A) The ATD-LBD angles (angle between a line passing through the COM of the R1 and R2 lobes of the ATD and a line passing through the COM of the D1 and D2 lobes of the LBD) for the seven structural classes show conformational differences between the two GluN2A subunits, breaking C2-symmetry. Side view of the GluN2A ECD for chains B and D of the 2-knuckle-asym triNMDAR conformation is shown to the right, with the LBD aligned. (B) The distance between the lower lobe of the ATD and the upper lobe of the LBD for GluN2A. An asymmetry exists between the ATD-LBD distances of the two GluN2A subunits, with this distance being consistently higher for chain D than chain B. This distance most dramatically changes for the E and E2 conformations. A side view of the two lobes for chains D of the 2-knuckle-sym and extended conformations is shown to the right, with the D1 lobes aligned. The black arrow indicates the direction of motion. (C) Inter-GluN2A LBD angles increase in the 1K-SS conformations as compared to the 2-knuckle conformations. and are the angles between the line passing through the COMs of the D2 lobes of GluN2A and lines passing through the COMs of the D1 and D2 lobes of chains B and D, respectively. Side view of the GluN2A LBD of the 2-knuckle-sym and 1-knuckle conformations is shown to the right, with the pseudosymmetry axes aligned.

We speculate that the GluN2A ATD clamshell of the subunit with the higher ATD-LBD angle undergoes clamshell closure first. In the 2-knuckle-asym conformation only the subunit with the higher ATD-LBD angle harbors a closed ATD clamshell, while the subunit with an open ATD clamshell retains the same ATD-LBD angle as the 2-knuckle-sym conformation. Even in the case of the triheteromeric receptor where in the ‘EDTA’ state GluN2A and GluN2A* subunits were both able to occupy the higher or lower ATD-LBD angle, with the addition of 1 μM zinc, the 2-knuckle-asym conformation only showed the GluN2A subunit occupying the higher ATD-LBD angle with a closed ATD clamshell. This observation is consistent with the order of the ATD clamshell closure being determined by GluN2A subunit’s ATD-LBD angle, as well as the GluN2A subunits in the receptor having the ability to switch readily between the high and low ATD-LBD states when both ATD clamshells are open.

The conformational changes at the ATD are efficiently transduced to the LBD for the 2- and 1-knuckle conformations. The ATD to LBD linker for GluN2A plays a central role in the transduction of the ATD allosteric modulation to the receptor gate (Gielen et al., 2009). The sequence of this linker region is highly divergent among the GluN2 subtypes. To understand the conformation and role of the ATD-LBD linker, we focused on the distance between the COM of the R2 lobe of the ATD to the D1 lobe of the LBD (Figure 5B). The R2-D1 COM distance does not change substantially (magnitude of change generally <0.6 Å) until the separation of the ATD dimer in the extended conformation, where the R2-D1 distance increases by ~6.1 Å. We speculate that the consistency of the R2-D1 distance in GluN2A allows the movements of the ATD R2 lobes to be directly translated to the LBD and not attenuated due to a flexible linker as is the case of AMPA receptors. The efficient transfer of ATD conformational changes to the LBD remains until the extended conformation, where the linker on chain D is stretched, and the ATD heterodimer is uncoupled from the receptor.

The lateral movement of the ATD R2 lobes at the pseudo-symmetry axis upon the closure of both ATD clamshells (Figure 4C) ‘pushes’ the opposing D1 lobes of GluN2A directly away from each other and brings the D2 lobes closer together at the LBD. To quantify the LBD movement, we connected the COM of the opposing D2 lobes of GluN2A with a vector. The angle that this vector makes with vectors passing through the COMs of D2 and D1 lobe of chains B and D are labeled and respectively (Figure 5C). Movement of the opposing D1 lobes away from each other while the D2 lobes come closer increases both δ and ε. The values for δ and ε remain indistinguishable when transitioning from the 2-knuckle-sym to the 2-knuckle-asym conformations, consistent with observations that the closure of a single ATD clamshell does not result in the lateral movement of R2-lobes. There is a ~7.4° increase in both δ and ε for the 1-knuckle conformation as compared to the 2-knuckle-sym conformation. The change in δ and ε from the 2-knuckle-sym conformation is slightly smaller for the extended conformation (~+6.5°) and even smaller in the extended-2 conformation (~+3.9°). The decrease in δ and ε when comparing the 1-knuckle state to the extended states is consistent with the less efficient transfer of the zinc-induced conformational changes at the ATD to the LBD for the extended conformations. In the splayed conformations, δ and ε increase by >30° as the splayed ATDs move the D1 lobes far away from each other.

### Channeling inhibition to the pore

How are changes at the ATD and LBD layers conveyed to the pore? Helix E in the D2 lobe of the LBD is directly connected to the M3 helix. The M3 helices, in turn, form the ion channel gate. We measured the distance between the E helices of the GluN2A subunits. We speculate that the larger the helix E separation, the greater the tension on the M3 helices and the higher probability of finding the gate in an open conformation. The rocking motion of LBDs upon zinc binding (Figure 5C) brings the D2 lobes closer together, resulting in the reduction of opposing helix E separation. This distance remains similar in the two 2-knuckle conformations and is reduced by 3.2 Å and 2.5 Å in the 1-knuckle and extended conformations, respectively (Figure 6A). The opposing helix E distance is similar in the 2-knuckle states and the extended-2 state, and this distance is reduced substantially (23 Å – 56 Å) in the splayed and super-splayed conformations. Consistent with the LBD rocking motion (Figure 5C), where the effects of zinc binding at the ATD was transferred to the LBD with lower efficiency for the extended and extended-2 conformations, the helix E separation in these conformations is also larger than the 1-knuckle conformation. The larger helix E separation for the extended and extended-2 conformations will likely result in a higher PO for these states as compared to the 1-knuckle state. The splayed open conformations show the lowest helix E separation, consistent with these classes having a PO value close to zero.

**Figure 6.**
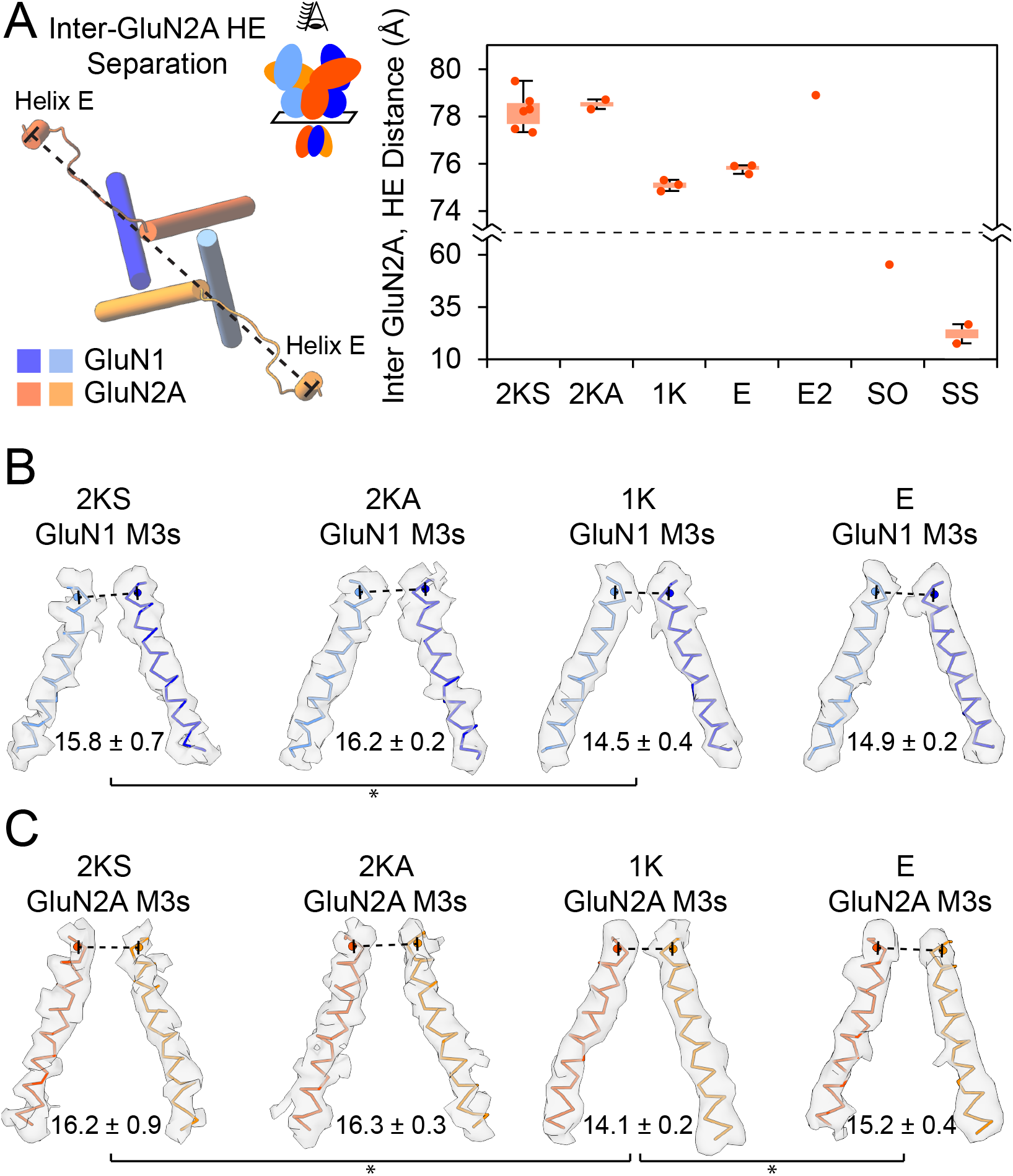
Measuring the Inhibition at the Pore. (A) Distance between opposing GluN2A Helix E (HE) COMs. Top view of the M3 helix bundle, GluN2A HEs, and the linkers which connect the two are shown to the right. (B and C) Side view of the M3 helix model of GluN1 (B) and GluN2A (C) with the associated density. Representative figures are from the highest resolution structures from each structural class. The diameter of the gate (distance between the COMs of residues 649-652 of GluN1 and 647-650 of GluN2A) is indicated by the dashed line. The value represents the mean and StDev for each structural class. * denotes significance (p < 0.05) using a two-tailed homoscedastic student’s t-tests.

Finally, we analyzed the conformational changes at the receptor gate itself. For this measurement, we focused only on the structures where the resolution was sufficient to rigid-body fit individual TMDs into the cryoEM maps. We measured the distance between the COM of residues 4-7 of the lurcher motif (SYTANLAAF) in GluN1 (Figure 6B) or GluN2A (Figure 6C) subunits (Zuo et al., 1997). We do not believe that any of our structures show the pore in the fully-open conformation, such as that found in molecular dynamics simulations (Song et al., 2018). However, there is a significant reduction in the level of pore opening between the 2-knuckle states and the 1-knuckle state in both GluN1 (1.3 Å, p = 0.02) and GluN2A (2.1 Å, p = 0.01) subunits. Consistent with the previous LBD and TMD observations, the M3 separation is similar between the two 2-knuckle states. The extended conformation shows a larger M3-M3 separation compared to the 1-knuckle conformation, with this difference being significant for the GluN2A subunits (1.1 Å increase, p = 0.02).

### Structural mechanisms for zinc and proton inhibition in NMDA receptors

We propose the following mechanism for zinc-inhibition of the GluN1/GluN2A NMDARs (Figure 7). The GluN1/GluN2A receptor in the absence of zinc and presence of agonists adopts a ‘2-knuckle-sym’ conformation, characterized by both GluN2A subunits in the receptor assuming the open ATD clamshell conformation. The R2 lobes of opposing GluN2A subunits are positioned back-to-back at the pseudo-symmetry axis. Upon the binding of zinc, the receptor occupies the 2-knuckle-asym, 1-knuckle, extended, extended-2, splayed-open and super-splayed conformations. When the receptor transitions from the 2-knuckle-sym to the 2-knuckle-asym conformation, the GluN2A subunit with the larger ATD-LBD angle undergoes ATD clamshell closure while the other ATD clamshell in the other subunit remains open. The closure of a single ATD clamshell is not sufficient to move the R2 lobes of GluN2A from their original back-to-back position at the pseudo-symmetry axis, and leads only to minor structural changes at the LBD and no change in the tension pulling on the gate. Closure of both ATD clamshells moves the R2-lobes of GluN2A away from each other at the pseudosymmetry axis and leads to the rocking motion of the GluN2A LBD clamshells, moving the opposing D1 lobes of the LBD away from each other and bringing the D2-lobes closer together, thus relieving tension in the LBD-TMD linkers. In the extended conformations, one ATD dimer loses interactions with its LBD and thus its modulation of receptor activity. Following further increases in protons and/or zinc, the receptor adopts splayed-open and super-splayed conformations where the LBD D1-D1 interface is disrupted, leading to a large rocking motion of the LBD dimers and releasing tension on the LBD-TMD linkers.

**Figure 7.**
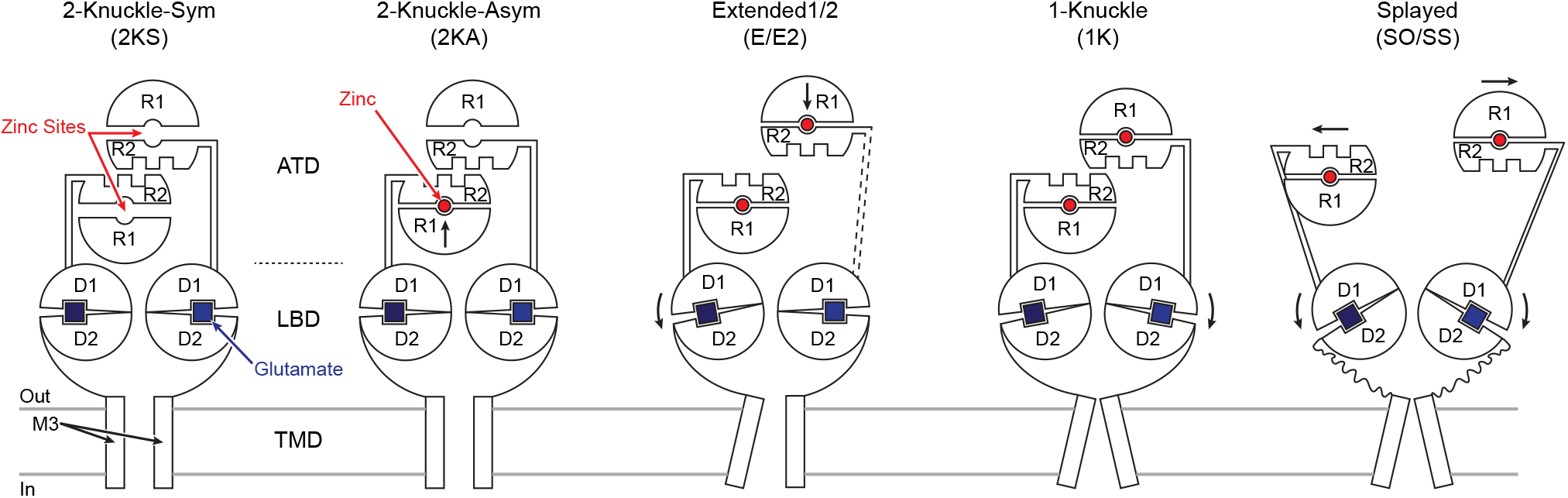
Schematic Summary for Activation and Zinc-Inhibition of GluN2A Containing NMDARs. Shown are the schematics for the conformational changes in GluN2A subunits which lead from active conformations (2KS/2KA) to lower open-probability states (E-E2) and finally to inhibited states (1K/SO/SS). These changes originate from the closure of both ATD clamshells in the presence of zinc, leading to the rocking of the LBD clamshells to bring their lower clobes closer together, and releasing the tension on the gate.

Based on the LBD rocking movements, helix E/helix E separation, and the gate diameter measurements, we speculate that the two 2-knuckle states are functionally equivalent, and represent the receptor in the highest P_O_ states. The extended and extended-2 conformations represent the next highest P_O_ states. Surprisingly, based on the LBD measurements, the extended-2 conformation should exhibit a similar P_O_ to the 2-knuckle states. The 1-knuckle and splayed conformations represent the lowest P_O_.

Our findings provide a structural justification for why we observe a higher degree of zinc-inhibition at low pH values, as we did not observe any classes with the 2-knuckle or extended-2 conformations under these conditions and all of the particles were in the lower P_O_ conformations. Similarly, we did not observe any classes other than the 2-knuckle conformations for the receptor at pH 8.0, representing higher P_O_. We can also explain why at 1 mM zinc, when we observe a large portion of the particles in the splayed conformations, we do not observe a larger degree of voltage-independent zinc-inhibition by electrophysiology (Paoletti et al., 1997). Even though at 1 mM zinc, ~40% of the particles are in the splayed conformations, ~60% of the particles are in the extended-2 conformation, representing high P_O_. Finally, we can account for the lower level of zinc-inhibition in the GluN1A/GluN2A/GluN2A* triNMDAR vs. the diNMDAR, as >50% of particles in the triNMDARs are in the 2-knuckle states vs. <20% for the diNMDARs. We were also able to demonstrate the reversibility of splayed and super-splayed conformation, and demonstrated that with the removal of zinc the receptor reverts back to the 2-knuckle-sym conformation.

The GluN1/GluN2A receptor adopts an ensemble of structural states upon binding zinc. As predicted, the receptor changes its occupancy among these structural states depending on zinc and proton concentrations (Amico-Ruvio et al., 2011; Dolino et al., 2017). The 1-knuckle conformation, a structure that the receptor adopts when inhibited by zinc and protons, resembles the GluN1/GluN2B receptor bound by ifenprodil or Ro25-6891 (Tajima et al., 2016; Zhu et al., 2016). The ‘splayed’ conformations, structural states seen in high concentrations of zinc or proton, display disrupted D1-D1 interfaces at the LBD dimer. Thus, we are able to reconcile seemingly contradictory findings in the field that zinc-inhibition in GluN2A receptors sometimes occurs through conserved structural mechanism of ifenprodil inhibition in GluN2B (Sirrieh et al., 2015), and at times through entirely different mechanisms involving the disruption of the D1-D1 interface (Gielen et al., 2008). Furthermore, through solving the structure of the GluN1/GluN2A/GluN2A* triheteromeric receptor in the presence of zinc, we showed that in some structural states both ATD clamshells were closed, demonstrating the propagation of the effects for a single ATD clamshell closure through the receptor.

### Experimental Procedures

More detailed experimental procedures are described in Supplemental Experimental Procedures.

#### Protein Construction, Expression, and Purification

DNA for the first 847 residues of the *rattus norvegicus* Grin1-1a (P35439-1) and the first 866 residues of Grin2A (G3V9C5) was cloned into Bacmam expression vectors (Goehring et al., 2014). Both genes contain an EGFP-tag on their C-terminus, with an 8XHIS-tag on the GluN1 vector and a Strep-tag II on the GluN2A vector, following EGFP. These constructs are referred to as N1_EM_ and N2A_EM_. For the triheteromeric receptor (triNMDAR), the N1_EM_ plasmid was modified by the removal of the 8XHIS-tag (N1_EMNoHIS_). A new version of N2A_EM_ was constructed with an 8XHIS-tag replacing the Strep-tag II (N2AEMHIS). Finally, another version of the N2AEM was constructed with an H128S mutation, as well as an N687Q mutation (N2A_EMZn-_). For radiolabeled functional assays, the N2A_EM_’s Strep-tag II was exchanged for a high-affinity streptavidin binding peptide tag (DNSA-tag, AA sequence: WIDFNRWHPQSGLPPPPILR, Takahashi et. al, manuscript in preparation). Baculovirus was generated in sf9 insect cells, and P2 virus was used to infect tsa201 cells at a multiplicity of infection (MOI) of 1:1. The receptors were purified as previously reported (Zhao et al., 2016; Zhu et al., 2016). In brief, the diNMDAR was purified using the Strep-tag II on the GluN2A subunit, and the triNMDAR was first purified using the Strep-tag II on the GluN2A* subunit, followed by HIS-tag purification on the wt-GluN2A subunit. Both receptors were then further purified by SEC. For the structural experiments of the receptor in the absence of zinc, EDTA was used to chelate any zinc contaminations introduced either during the purification of the receptor, or during the blotting step of the cryoEM sample preparation.

#### Two Electrode Voltage Clamp (TEVC) Electrophysiology

The N1_FM_ and N2A_FM_ constructs were subcloned in pGEM vector and *in vitro* transcribed to generate mRNAs. *Xenopus laevis* oocytes were injected with 0.5 ng total mRNA (equal amounts of N1 and N2A) and after 12 – 24 hrs of incubation at 18 °C currents were measured using two-electrode voltage clamp (Axoclamp 200B) at a holding voltage of −60 mV. The base recording solution was (in mM): 60 NaCl, 40 HEPES, 10 Tricine, buffered initially to pH 10.3 and then re-buffered to different pH using HCl. Zinc was added from a 100 mM stock of ZnCl_2_ and free-zinc was estimated as: [Zn]_total_/200 (Gielen et al., 2008). Recordings were carried out in triplicates. Data was analyzed using MATLAB.

#### Scintillation Proximity Assay (SPA)

SPA assays were set up using 40 nM purified protein in 1X HBS (pH 7.4) supplemented with 0.1% digitonin, 10 mM tricine, 0.5 mg/mL of YSi-streptavidin beads, and 100 nM [^3^H]MK-801 per well. Various additives such as 100 μM glutamate/glycine, and ZnCl_2_ were added to the wells, but the final volume was kept at 100 μL. Samples were run in triplicates. Concentration of free zinc was estimated as described above. The plate was placed inside a microbeta TriLux 1450 LSC plate reader and read every 12 minutes. After allowing 12 hours for [^3^H]MK-801 to reach equilibrium, 100 μM nonlabeled MK-801 was added to each sample to prevent the re-binding of radiolabeled ligand, and the plate was placed back in the plate reader to measure the off-rate of MK-801. Data was analyzed using a one-site binding model.

#### Electron Microscopy Sample Preparation and Image Acquisition

SEC purified protein was used for cryoEM sample preparation. QUANTIFOIL AU 1.2/1.3 300 mesh grids were glow discharged for 1 minute at 15 mA with the carbon side facing up. For all conditions, 1 mM glutamate and glycine were added to the samples. For low zinc concentration samples (1 μM), concentrated protein was placed in a dialysis bag (heavy-metal free Dispodialyzer) and dialyzed against 2000x volume of a buffered solution containing 0.1% digitonin, 1 mM glutamate/glycine and 1 μM ZnCl_2_. The sample was dialyzed for a minimum of 4 hours and then used for grid preparation. Vitreous sample was prepared by applying 3-4 μL of protein solution at ~4 mg/mL, blotting the excess solution away for 3 seconds using the FEI Vitrobot at 18 °C and 100% humidity, and plunge-freezing the grid in liquid ethane.

For all of the datasets except for the diheteromeric receptor at pH 7.4 with 1 mM EDTA, data collection was carried out at Oregon Health and Science University Multiscale Microscopy Core using a 300 keV FEI Titan Krios equipped with an energy filter (Gatan Image Filter). Images were collected using a Gatan K2 direct electron detector in superresolution mode at 81 kx magnification (unbinned pixel size of 0.855 Å/pixel). Images were collected using serialEM (Mastronarde, 2005) with a typical defocus range of −1 – – 2.5 μm and with a total dose of ~55 e^-^/Å^2^. The diheteromeric receptor at pH 7.4 with 1 mM EDTA dataset was collected at the Howard Hughes Medical Institute’s CryoEM Facility at the Janelia Research Campus, using an FEI Titan Krios equipped with a Cs corrector and an energy filter (Gatan Image Filter). This dataset was collected using the super resolution mode on a K2 direct electron detector in super-resolution mode with an unbinned pixel size of 0.66 Å/pixel.

#### Micrograph Processing, 3D-Classification and Refinement

Image stacks were aligned to compensate for motion during recording using MotionCor2 (Zheng et al., 2017), Fourier space binned 2 × 2, and the contrast transfer function was estimated using Gctf (Zhang, 2016). Template-free automatic particle selection was done using DoG-Picker (Voss et al., 2009) or gAutomatch (Unpublished: URL http://www.mrc-lmb.cam.ac.uk/kzhang/Gautomatch/). Reference-free 2D-classification of particles was carried out in RELION (Fernandez-Leiro and Scheres, 2017; Scheres, 2012) followed by manual inspection, selection of the classes representing NMDARs, and further classification. Generally, three rounds of 2D-classification were performed with binned particles (initial round with particles binned by 5, subsequently binned by 3, followed by the last round without binning). After each round, particles were re-centered and re-extracted with the new binning factor. After 2D-cleanup of reference free picked particles, ~30 of the 2D class averages, representing a diverse set of projections, were used for template-based particle picking by gAutomatch. This set of particles was also cleaned up through three rounds of 2D-classification as described above. The two cleaned up particle sets (template-free and template-based) were then combined, and duplicate particles were removed by eliminating any particle whose center was closer than 90 Å to another particle. This combined dataset was then used in 3D-classification and refinement.

The initial model for classification was *de novo* generated using CryoSPARC (Punjani et al., 2017). This model was then used in RELION for 3D-classification. Focused classification (focusing on ECD) was used in the datasets to better separate classes with similar structural features. Some datasets were classified into different number of classes (i.e. 9, 7, and 5), and the classes with the same structural conformations were combined, and duplicate particles removed. After additional rounds of refinement/further classification, the final particle set for each class was exported to cisTEM (Grant et al., 2018). Further refinement in cisTEM was carried out by masking out the micelle, while keeping the TMD of the receptors for alignment. The area outside of the mask was filtered to 20 Å and down-weighted for alignment. For a more accurate and conservative resolution estimate, the cisTEM-generated half-maps were used in RELION to generate the Fourier Shell Coefficient (FSC) curve, and the resolution was calculated at FSC = 0.143 (Scheres and Chen, 2012). Local resolution estimations were generated using Bsoft (Heymann et al., 2008). For a higher resolution reconstruction of the 2 knuckle asym class of the TriNMDAR, a focused refinement based on an ECD mask was performed. The resolution of the final reconstruction was improved by ~0.5 Å. The final maps were sharpened by Phenix. Autosharpen (V1.13) using the resolution of the map obtained from cisTEM (Adams et al., 2010; DiMaio et al., 2013). The splayed open and super splayed open classes were not refined in cisTEM, and the maps from those classes, as well as their FSC curves, were generated by RELION.

#### Model Generation, Fitting, Refinement, and Structural Analysis

The GluN1 homology model was constructed by SWISS-MODEL (Arnold et al., 2006) based on the crystal structure of the GluN1/GluN2B (PDB code: 4PE5) (Karakas and Furukawa, 2014). The GluN2A homology model was based on three sources. The ATD of the GluN2A homology model was based on the crystal structure of the isolated GluN1/GluN2A ATD (Romero-Hernandez et al., 2016) (PDB code: 5TQ0), the D1 lobe of the LBD was based on the crystal structure of the isolated LBD of GluN1/GluN2A (Yi et al., 2016) (PDB code: 5I57), and the remainder of the homology model was based on the cryoEM structure of the GluN1/GluN2A/GluN2B triheteromeric NMDAR (Lu et al., 2017) (PDB code: 5UOW). Each subunit was split into 5 parts (R1, R2, D1, D2, and TMD). The 5 pieces of each subunit (20 in total) were independently rigid-body fitted into each of the cryoEM density maps using UCSF Chimera (Pettersen et al., 2004). The only exceptions to this procedure were the splayed open structure, for which the TMD of the channel was fitted as a single rigid-body, and the super splayed structures, where each subunit’s domain was fitted (instead of each lobe) and the TMD tetramer was fitted as a single rigid-body. Each model was then manually inspected and corrected in COOT (Emsley and Cowtan, 2004), followed by real-space refinement in Phenix (Adams et al., 2010). All center of mass calculations are based on the center of mass of the alpha-carbon of the noted residues (Figure S8). In the case of the ATD clamshell closure angles, the alpha carbon of Ser 150 residue was used as the pivot point along with the COMs of R1 and R2 lobes of GluN2A to measure the clamshell angle. FSC curves correlating the final models to the cryoEM maps were generated in Phenix. All the figures were made using UCSF’s Chimera and PyMOL (Yuan et al., 2016).

## Supplemental Information

Supplemental Information includes Supplemental Experimental Procedures, eight figures, and four tables.

## Author Contribution

Conceptualization, F.J.Y., S.C., and E.G.; Methodology, F.J.Y. and S.C.; Investigation, F.J.Y. and S.C.; Formal Analysis and Visualization, F.J.Y.; Writing – Original Draft, F.J.Y.; Writing – Review & Editing, F.J.Y. and E.G.; Supervision, E.G.; Funding Acquisition, F.J.Y., S.C., and E.G.

## Acknowledgements

We thank Z.H. Yu, H.T. Chou, and R. Huong from Janelia Research Campus for assistance with data collection. Electron microscopy was performed at Oregon Health and Science University (OHSU) at the Multiscale Microscopy Core (MMC) with technical support from the Oregon Health & Science University (OHSU)-FEI Living Lab and the OHSU Center for Spatial Systems Biomedicine (OCSSB). Elemental analysis to quantitate background zinc contamination was performed in the OHSU Elemental Analysis Core. We thank all of the members of the Gouaux, Baconguis, and Whorton labs for helpful discussions. We thank A.J. Romero for help with the illustrations, and we thank H. Owen, N. Yoder, and V. Navratna for proofreading. F.J.Y is supported by the National Institutes of Health (1F32MH115595). S.C. is supported by the Jane Coffin Childs Fund (61-1591). This work was supported by the National Institutes of Health (R01NS038631). E.G. is an investigator with the Howard Hughes Medical Institute.

## Supplemental Information titles and legends

**Figure S1. CryoEM Maps and Properties of GluN2A containing NMDARs in the Absence of Zinc at pH 7.4**

(A-C) Representative cryoEM micrographs for diNMDAR (A) or the triNMDAR (B) with 1 mM EDTA, or diNMDAR with 1 mM ZnCl_2_ and 3 mM EDTA (C) at pH 7.4. (D, G, K) Final cryoEM maps of the diNMDAR (D) or triNMDAR (G) in the presence of 1 mM EDTA, or the diNMDAR in the presence of 1 mM ZnCl_2_ and 3 mM EDTA (K), colored based on the local resolution estimation by Bsoft. (E, H, L) Euler angle distribution of particles in the cryoEM maps. (F, I, M) the Fourier shell correlation curves for the cryoEM maps without masking (cyan), or by providing a soft solvent-mask (black). The FSC curve between the atomic model and the map is shown in green. (J) Side views (left) of the diNMDAR cryoEM map in the presence of agonists (1 mM glutamate and glycine), 1 mM ZnCl_2_, and 3 mM EDTA. Top view of the atomic model for the ATD is shown on the right. The rounded rectangles represent the ‘knuckles’ and show the interaction between the α5 and α6 helices of opposing GluN2A subunits.

**Figure S2. CryoEM Maps and Properties of GluN1/GluN2A diNMDAR in 1 μM zinc at pH 7.4**

(A) Representative cryoEM micrograph for the diNMDAR in 1 μM zinc at pH 7.4 (B) Side view of the ECD (top) and top view of the ATD (bottom) for the 3D classes in this condition, prior to final refinements (C, F, I) Final cryoEM maps of the 2-knuckle-asym (C), 1-knuckle (F), and extended (I) conformations in the presence of 1 μM ZnCl_2_, colored based on the local resolution estimation by Bsoft. (D, G, J) Euler angle distribution of particles in the cryoEM maps. (E, H, K) the Fourier shell correlation curves for the cryoEM maps without masking (cyan), or by providing a soft solvent-mask (black). The FSC curve between the atomic model and the map is shown in green.

**Figure S3. CryoEM Maps and Properties of GluN1/GluN2A/GluN2A* triNMDAR in 1 μM zinc at pH 7.4**

(A) Representative cryoEM micrograph for the triNMDAR in 1 μM zinc at pH 7.4 (B) Side view of the ECD (top) and top view of the ATD (bottom) for the 3D classes in this condition, prior to final refinements (C, F, I, L) Final cryoEM maps of the 2-knuckle-sym (C), 2-knuckle-asym (F), 1-knuckle (I), and extended (L) conformations in the presence of 1 μM ZnCl_2_, colored based on the local resolution estimation by Bsoft. For the 2-knuckle-asym (F) the figures at the bottom show the side-views of the receptor when the refinement was confined to the ECD portion of the receptor only. (D, G, J, M) Euler angle distribution of particles in the cryoEM maps. (E, H, K, N) the Fourier shell correlation curves for the cryoEM maps without masking (cyan), or by providing a soft solvent-mask (black). The FSC curve between the atomic model and the map is shown in green. For the 2-knuckle-asym conformation (H), the grey and light green lines represent the cryoEM map FSC and map-to-model correlation FSC for the ECD based refinement, respectively.

**Figure S4. CryoEM Maps and Properties of GluN1/GluN2A diNMDAR in 1 μM zinc at pH 6.1**

(A) Representative cryoEM micrograph for the diNMDAR in 1 μM zinc at pH 6.1 (B) Side view of the receptor (top) and top view of the ATD (bottom) for the 3D classes in this condition, prior to final refinements (C and F) Final cryoEM maps of the 1-knuckle (C) and extended (F) conformations in the presence of 1 μM ZnCl_2_, colored based on the local resolution estimation by Bsoft. (D, G, I) Euler angle distribution of particles in the cryoEM maps. (I) represents the Euler angle distribution of particles for the super-splayed conformation. (E, H, J) the Fourier shell correlation curves for the cryoEM maps without masking (cyan), or by providing a soft solvent-mask (black). The FSC curve between the atomic model and the map is shown in green.

**Figure S5. CryoEM Maps and Properties of GluN1/GluN2A diNMDAR in 1 mM zinc at pH 7.4**

(A) Representative cryoEM micrograph for the diNMDAR in 1 mM zinc at pH 7.4 (B) Side view of the receptor (top) and top view of the ATD (bottom) for the 3D classes in this condition, prior to final refinements (C and F) Final cryoEM maps of the extended-2 (C) and splayed-open (F) conformations in the presence of 1 mM ZnCl_2_, colored based on the local resolution estimation by Bsoft. (D, G, I) Euler angle distribution of particles in the cryoEM maps. (I) represents the Euler angle distribution of particles for the super-splayed conformation. (E, H, J) the Fourier shell correlation curves for the cryoEM maps without masking (cyan), or by providing a soft solvent-mask (black). The FSC curve between the atomic model and the map is shown in green.

**Figure S6. CryoEM Maps and Properties of GluN1/GluN2A diNMDAR at pH 8.0**

(A and F) Representative cryoEM micrographs for the diNMDAR in 1 mM zinc at pH 7.4 (B and G) Side views (left) of the diNMDAR cryoEM map in the presence of agonists (1 mM glutamate and glycine), 0.1 mM EDTA (B) or 1 μM ZnCl_2_. Top view of the atomic model for the ATD is shown on the right. The rounded rectangles represent the ‘knuckles’ and show the interaction between the 5 and 6 helices of opposing GluN2A subunits. (C and H) Final cryoEM maps of the receptor in the presence of 0.1 mM EDTA (C) 1 μM zinc (F), colored based on the local resolution estimation by Bsoft. (D, I) Euler angle distribution of particles in the cryoEM maps. (E, J) the Fourier shell correlation curves for the cryoEM maps without masking (cyan), or by providing a soft solvent-mask (black). The FSC curve between the atomic model and the map is shown in green.

**Figure S7. TEVC traces for the diNMDAR and Representative Densities for 2-Knuckle-Asym triNMDAR**

(A) Representative zinc-inhibition TEVC trace for the GluN1/GluN2A cryoEM construct at pH 7.4 and 100 μM glutamate and glycine. (B) Extent of zinc-and proton-inhibition for the GluN1/GluN2A cryoEM diNMDAR constructs. (C-L) Densities for selected α-helices in GluN1 and GluN2A subunits of the 2-knuckle-asym conformation (ECD aligned map) and the associated molecular model.

**Figure S7. Sequence Homology and Predicted Secondary Structure of GluN1, GluN2A, and GluN2B.**

Predicted glycosylation sites for which we found corresponding density are marked in red and with the red arrow and the ‘Glyco’ label. The residues comprising each ECD lobe (used for COM calculations) are marked below the sequence. The GluN2A* mutations (H128S and N687Q) are marked in blue.

